# Assembly properties of FtsZ from cyanobacterium *Synechocystis* sp. PCC 6803

**DOI:** 10.1101/677963

**Authors:** Na Wang, Li Bian, Xueqin Ma, Yufeng Meng, Cyndi S. Chen, Mujeeb ur Rahman, Tingting Zhang, Zhe Li, Ping Wang, Yaodong Chen

## Abstract

Tubulin homologue FtsZ is the major cytoskeletal protein in the bacterial cell division machinery. Here, we studied the biochemical and assembly properties of SyFtsZ, FtsZ from cyanobacterium *Synechocystis* sp. PCC 6803. SyFtsZ had a slow GTPase activity of around 0.4 GTP per FtsZ per minute and assembled into thick, straight protofilament bundles and curved bundles designated toroids. The assembly of SyFtsZ in the presence of GTP occurred in two stages. The first stage was assembled into single straight protofilaments and opened circles; the second stage was association of the protofilaments into straight protofilament bundles and toroids. In addition to these assemblies in GTP, highly curved oligomers and minirings could be observed after GTP hydrolysis or in the presence of GDP. Those three types of protofilaments of SyFtsZ provide support for the hypothesis for a constriction force based on curved protofilaments.

---

FtsZ, a tubulin homologue, is a crucial protein in bacterial cell division and is well conserved in almost all bacteria, archaea and chloroplasts. It assembles spontaneously into single protofilaments (pfs), sheets and bundles *in vitro*, and it accumulates *in vivo* at the site of division early in the cell division process to form a dynamic protein complex, the contractile ring or Z-ring (1–3).

FtsZ from *Escherichia coli* (EcFtsZ) assembles *in vitro* into most single pfs with an average length of 200 nm (4). EcFtsZ pfs are highly dynamic, turning over with a half time of 5-8 seconds *in vitro* (5,6) and *in vivo* (7). The mechanism of dynamics is now established to be treadmilling both *in vitro* (8,9) and *in vivo* (10,11). Treadmilling involves small patches of membrane-bound pfs adding subunits at one end and disassembling at the other.

FtsZ from different species can have different dynamic properties. The Z-ring in *Bacillus subtilis* has a similar dynamic turn-over rate as in *E. coli* (7); however, the Z-ring from the slow-growing pathogen *Mycobacterium tuberculosis* has a slower turnover rate of 25 s half time (12). Chloroplast Z-rings have unique and much slower dynamic properties (13–15). Chloroplasts have two copies of FtsZ, designated FtsZ1 and FtsZ2 from green algae and plants, and FtsZA and FtsZB from red algae. Both FtsZs co-localize in the middle of the chloroplast and are required for division (16–18). *In vitro*, they assemble into thick straight bundles of pfs, either separately or as a mixture (19,20). Previous results suggested that one FtsZ might assemble as the primary scaffold and another co-assembly enhance polymers disassembly (20). The slow dynamic of FtsZ bundles might be related with the large diameter of the chloroplasts.

The chloroplast originated from a cyanobacterial endosymbiont, and had maintained FtsZ for its division. In contrast to the two FtsZs in chloroplasts, there is only one copy of FtsZ in cyanobacteria, similar to other bacteria. It should be interesting to compare the biochemical and assembly properties of cyanobacterial FtsZ to FtsZ from chloroplasts and bacteria. Here, we studied the biochemical properties of FtsZ from the cyanobacterium *Synechocystis* sp. PCC 6803 (SyFtsZ).

## RESULTS

### SyFtsZ assembles into straight bundles of pfs and toroids-like circle bundles

Negative stain electron microscopy (EM) images of SyFtsZ filaments assembled in GTP showed some isolated pfs, but mostly large bundles of pfs that were either straight or curved into toroids (Fig 1). The diameter of the toroids ranged from 200 to 300 nm. Straight bundles and toroids were mostly isolated, but occasionally, a transition between straight and curved bundles was observed (Fig 1, arrowed). In contrast to EcFtsZ, which can assemble in the absence of Mg^2+^ (6), SyFtsZ showed no assembly without Mg^2+^ or when using GMPCPP to replace GTP (data not shown).

**Figure 1.**
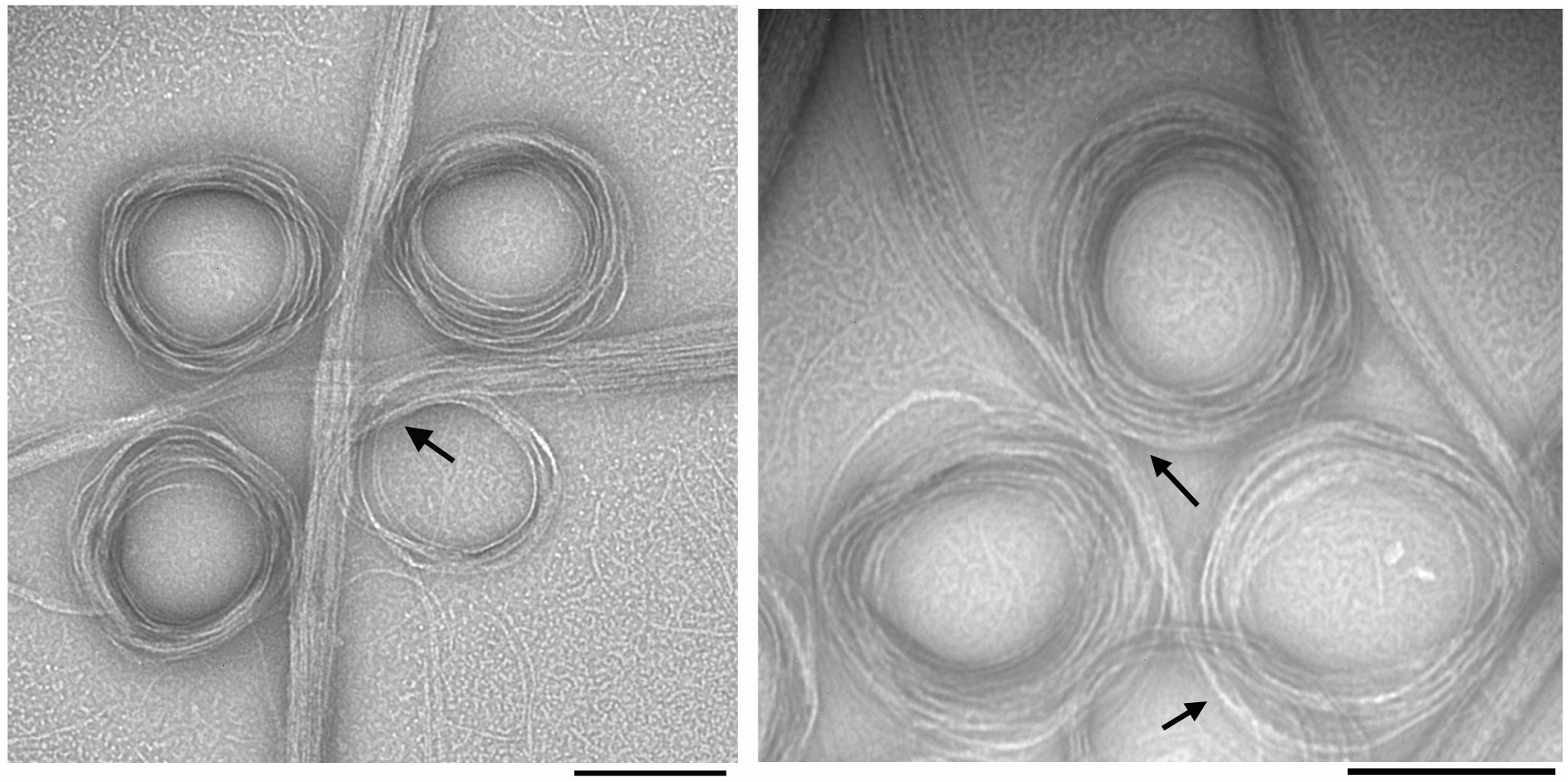
Electron microscopy (EM) images of negatively stained SyFtsZ pfs after adding 0.5 mM GTP. 10 μM SyFtsZ assembled into pfs (some isolated pfs are seen in the background), which further associate into straight bundles and toroids. The diameter of the toroids is 200-300 nm. Transitions between straight bundles and toroids are indicated by arrows. Bar is 200 nm.

Assembly was then monitored by light-scattering, which showed that it proceeded in two stages (Fig. 2). The light-scattering reached a final plateau at ~300 s (Fig. 2A). Amplification of the first 100 s (Fig. 2B) showed an early stage where a 15-20 s lag was followed by a rise to an intermediate plateau at 30-50 s. After 50 s the second stage began as light-scattering accelerated leading to the final plateau. Negative stain EM showed that the first stage resulted in the assembly of isolated short pfs and thin bundles of pfs curved into partial circles (Fig. 2C). The curved filaments consisted of apparently more than single filament, and their diameter were 200 to 300 nm, similar to toroids-like circle bundles, but mostly too short to form a complete circle. At the final plateau following the second stage of assembly, the EM showed predominantly large bundles of pfs and toroids.

**Figure 2.**
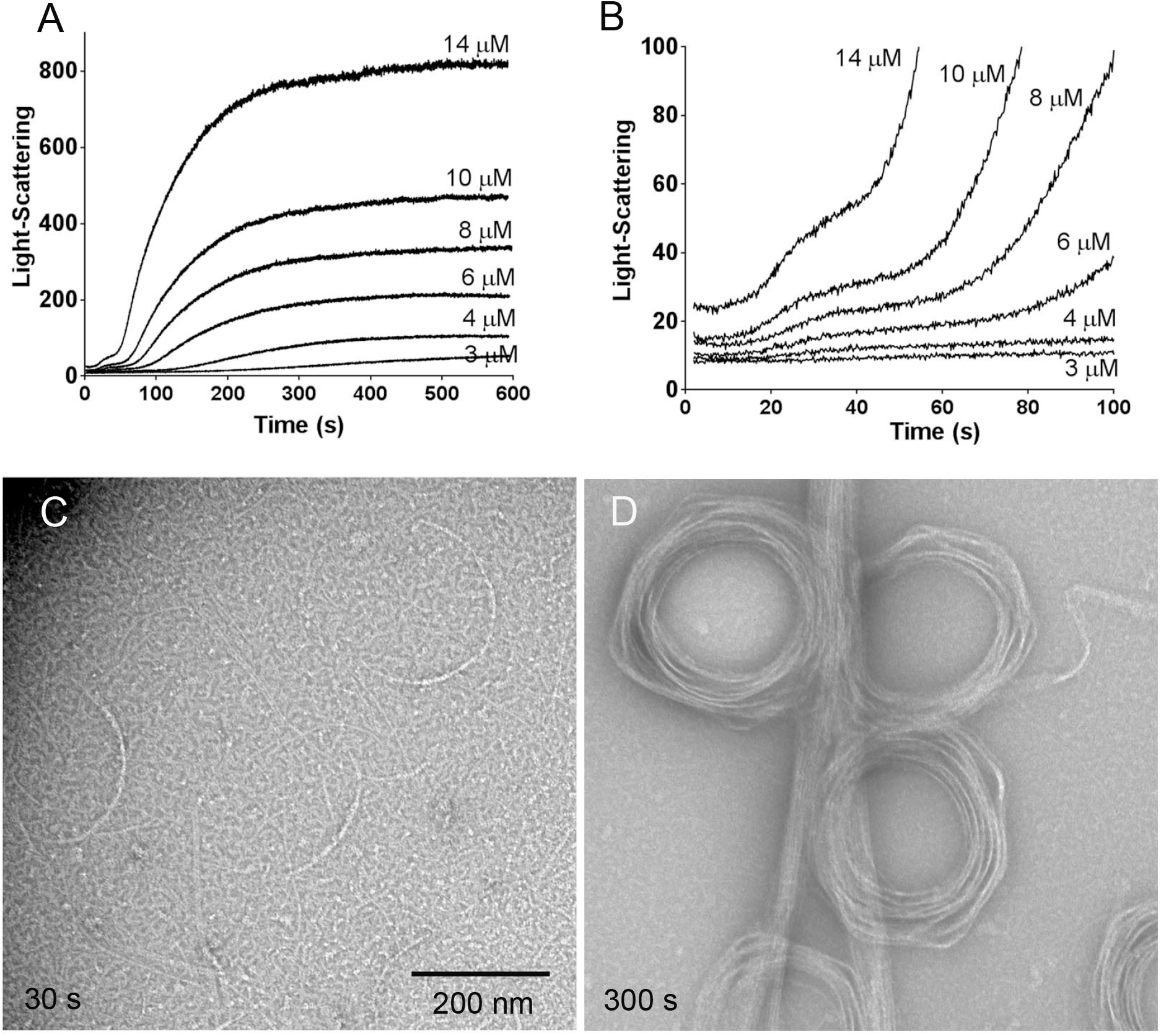
The assembly of SyFtsZ assayed by light-scattering shows two stages (A-B). Panel A shows that light-scattering reaches a final plateau at 300-600 s. Panel B amplifies the first 100 s and shows that light-scattering first approaches an intermediate plateau at 30-50 s and then accelerates to the much higher final plateau. EM of 10 μM SyFtsZ shows short single pfs and partial circles at the first stage assembly, 30 s (C), and assembly into straight bundles and toroids at the second stage assembly, 300 s minutes (D). Bar is 200 nm.

We repeated the assembly with a limited amount of GTP (0.1 mM GTP) and found that the light-scattering signal rose to a plateau at 500 seconds and then slowly dropped back to the base after 1 hour (Fig. 3A). This suggested that the large pf bundles disassembled after the GTP was used up. Fig. 3B shows an assay of GTPase as a function of FtsZ concentration. GTPase activity was 0.41 ± 0.2 GTP per FtsZ per minute above a critical concentration around 1.6 μM (Fig. 3B). This is approximately 10 times less than EcFtsZ, but close to that of chloroplast FtsZ. At this rate of hydrolysis, the 10 μM SyFtsZ in Fig. 2A should consume the 100 μM GTP in about 30 min. However, the polymers in that experiment started to disassemble after 10 min, well before the GTP should be exhausted. Small and Addinall (21) documented for EcFtsZ that GDP inhibited assembly, suggesting that the early decrease in light-scattering seen here was due to inhibition by the buildup of GDP.

**Figure 3.**
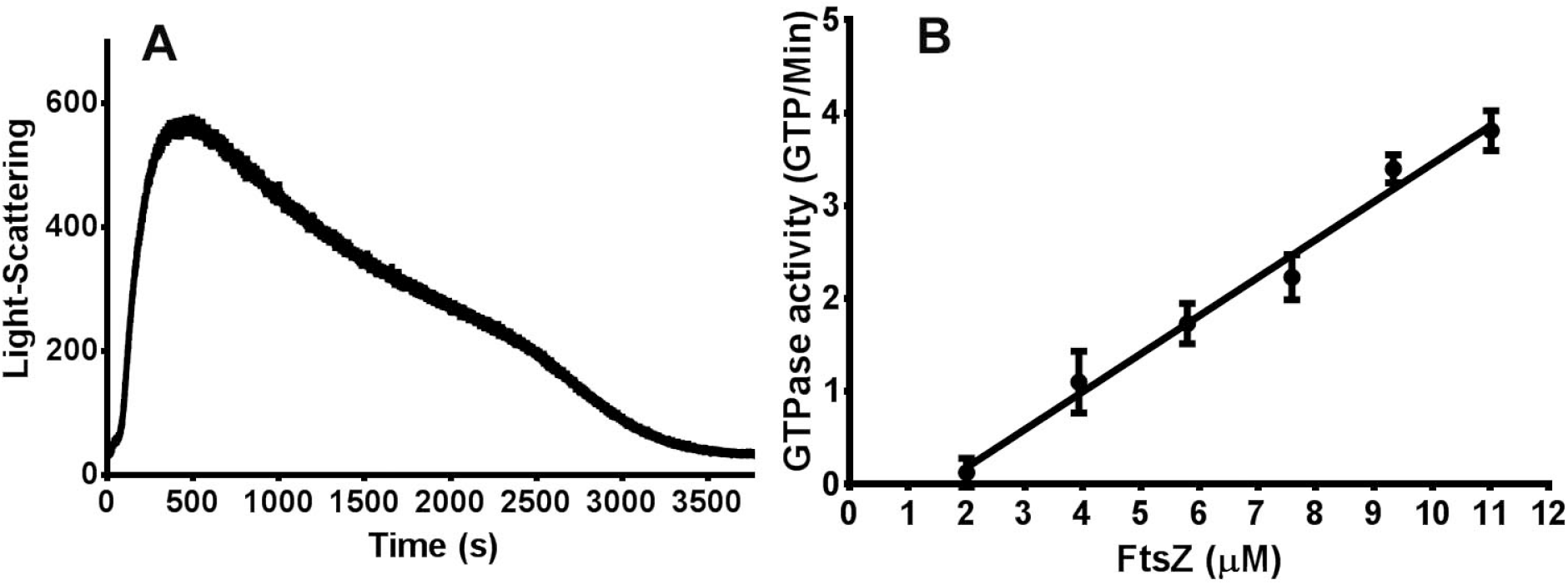
(A) Light-scattering assay of 10 μM SyFtsZ assembled in 0.1 mM GTP reached a plateau at 500 s, and then slowly decreased to the base after 1 hour. (B) The GTPase activity of SyFtsZ in HMK buffer was 0.41 ± 0.2 GTP per minute per FtsZ above a critical concentration 1.6 μM.

### Assembly properties of SyFtsZ change with pH

Next, we examined the assembly of SyFtsZ in buffers of different pH. The light-scattering assay showed two effects of lowering pH: an accelerated assembly and increased plateau (Fig. 4A). The assemblies displayed typical cooperative assembly kinetics with a lag time. We could not resolve two-stage assemblies at pH 7.2 and pH 6.5 buffers, perhaps because the first stage was too short or too weak. EM of the pH 7.2 assembly at 40 seconds showed short straight bundles and toroids already assembled, mixed with short pfs and partial circles (data not shown). EM of assemblies at the final plateau showed a mixture of straight protofilament (pf) bundles and toroids at pH 7.5, pH 7.2 and pH 6.8 (Fig. 4B-D). However, only straight bundles were observed at pH 6.5 (Fig. 4E). The increased light-scattering at reduced pH is consistent with the bundles being larger in diameter and/or longer.

**Figure 4.**
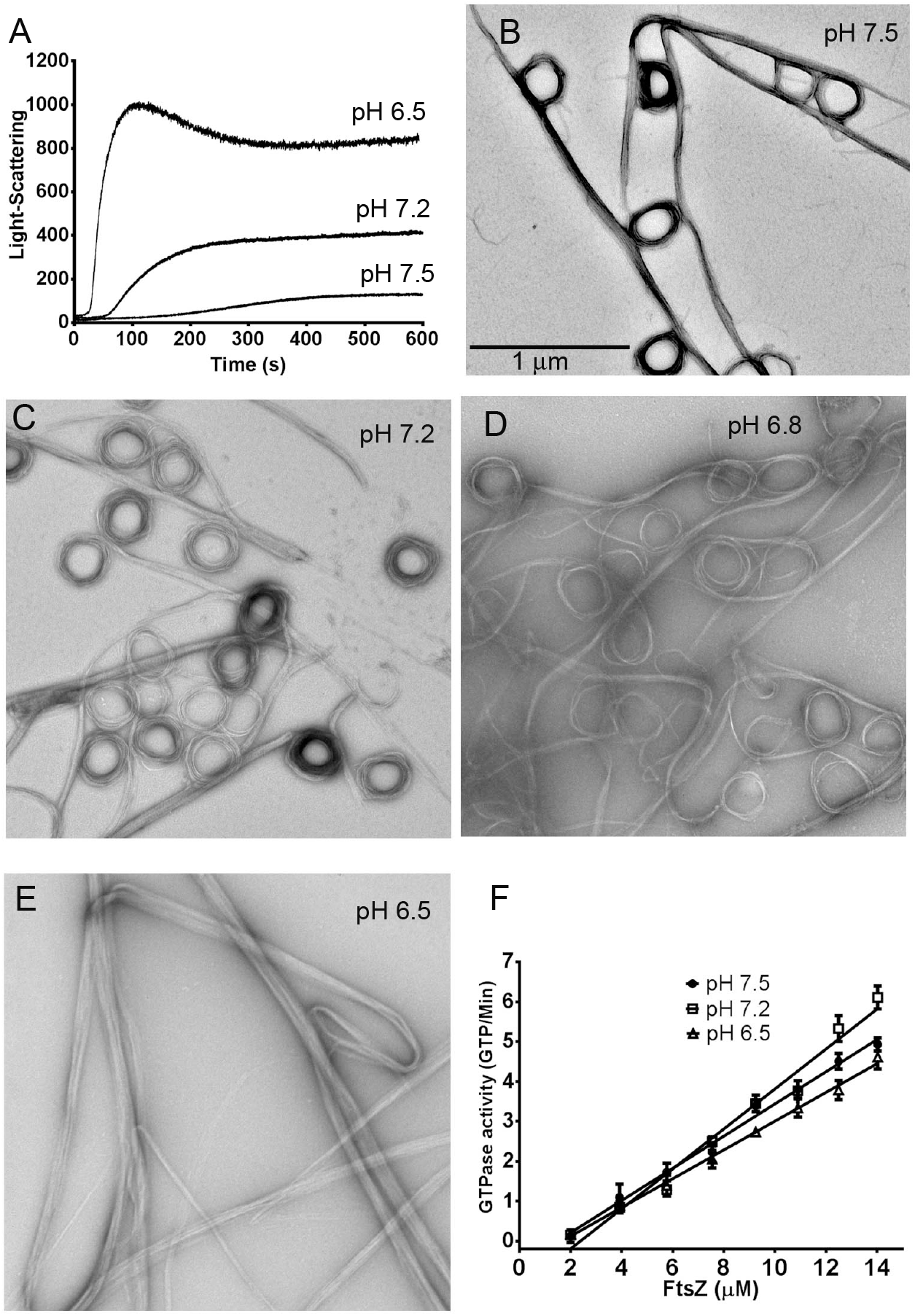
SyFtsZ assembly in buffers of different pH. (A) Assembly kinetics of 5 μM SyFtsZ monitored by light-scattering. The light-scattering signals are much higher when the pH dropped from 7.5 to 6.5. (B-E) Negative stain EM images of 10 μM SyFtsZ assembled at different pH, from 7.5 to 6.5. Bar is 1 μm. (F) GTPase activity of SyFtsZ at different pH show little change. All buffers contained 100 mM KCl, 5 mM MgCl2, and 0.5 mM GTP. pH 7.5 and 7.2 were buffered with HEPES; pH 6.8 and 6.5 were buffered with MES.

In contrast to the large changes in light-scattering and bundle morphology at lower pH, the GTPase was hardly changed. The GTPase activity of FtsZ was 0.41 ± 0.2 GTP per FtsZ per minute at pH 7.5, 0.50 ± 0.1 at pH 7.2 and 0.36 ± 0.2 at pH 6.5 with critical concentrations of 1.5-2 μM (Fig. 4F).

### SyFtsZ assembled into highly curved oligomers and miniring structures in the presence of GDP

Previous studies have shown that FtsZ from *E. coli* and *Pseudomonas aeruginosa* could assemble into highly curved short oligomers or miniring structures with the average diameter of 23 nm in the presence of GDP (22–25). Minirings were especially favored in the presence of ZipA protein (24). We also found highly curved oligomers and miniring structures in the EM images of SyFtsZ pfs during GTP hydrolysis (Fig. 5A) or in the presence of GDP (Fig. 5B,C) in both pH 7.5 and pH 6.5 solutions. The diameters of these highly curved pfs or minirings were ~20 ± 2 nm. Sedimentation results confirmed assembly of these oligomers. Using a Beckman TLA-100 rotor, we selected two centrifuge speeds: 50,000 rpm and 90,000 rpm. Only large bundles assembled in GTP could be spun down at 50,000 rpm (around 100,000 g); in contrast, both bundles in GTP and partly short oligomers and/or minirings could be spun down at 90,000 rpm (around 310,000 g). Figure 5 (D and E) show the sedimentation results analyzed by SDS-PAGE. At 50,000 rpm, there was little protein in pellet without GTP. At 90,000 rpm without GTP 21% of FtsZ was pelleted at pH 7.5 at both 5 μM and 10 μM FtsZ, and 49% of 10 μM FtsZ was pelleted at pH 6.5. The pellets are consistent with the short oligomers and minirings observed by EM.

**Figure 5.**
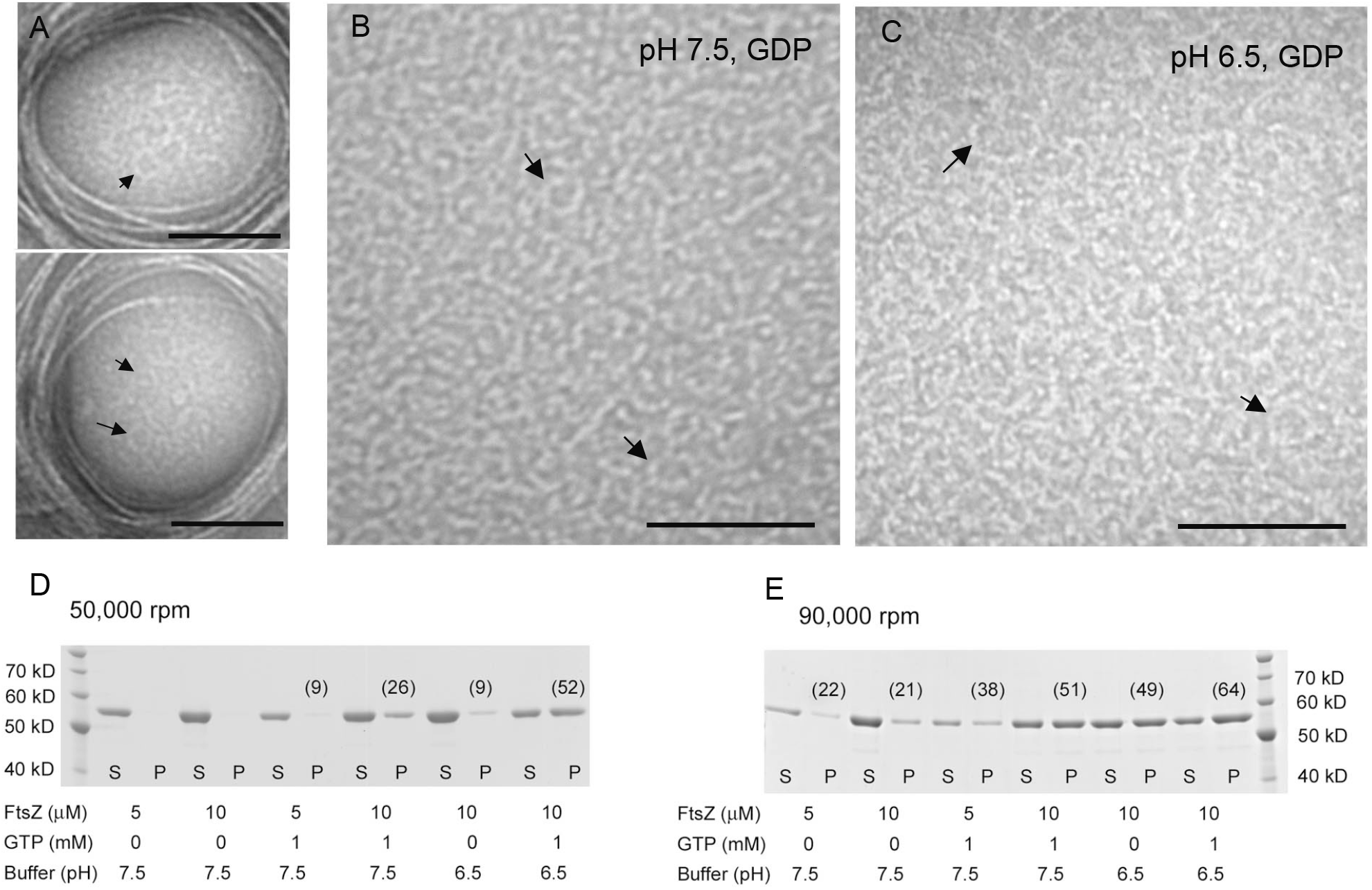
(A-C) SyFtsZ minirings observed by negative stain EM. (A) 10 μM SyFtsZ assembled in the presence of 0.5 mM GTP. In addition to straight bundles and toroids, some minirings are also observed. The diameter of miniring is ~20 nm. Bar is 100 nm. (B-C) 10 μM SyFtsZ assembled into oligomers and minirings in the presence of 0.5 mM GDP at pH 7.5 (B) or at pH 6.5 (C). Bar is 100 nm. Arrows point to minirings. (D-E) SDS-PAGE analysis of sedimentation of 5 μM or 10 μM SyFtsZ with or without GTP and centrifuged at 50,000 rpm (D) or 90,000 rpm (E) for 30 min in a Beckman TLA100 rotor. Numbers in parentheses indicate the percent protein in the pellet.

### Mutant A..A to T..S in the H8 helix reduces bundle formation and enhances GTPase activity

FtsZ comprises two globular subdomains, which are connected by a central core helix, H7 helix, and a synergy loop, T7 loop. The T7 loop is located between the H7 helix and H8 helix, and it reaches into the active site of an adjacent subunit to activate its GTPase activity (26–28)(Fig. 6A). The amino acids N207, D209 and D201 of EcFtsZ are highly conserved in all FtsZs. Mutation of D209 in the T7 loop or D212 in the H8 helix of EcFtsZ severely impaired its GTPase activity and function (29,30). Following the very conserved sequence DFADVR(K), we have identified two amino acids in the H8 helix that are conserved as Thr and Ser in fast-growing bacteria (T215 and S218 of EcFtsZ, or T216 and S219 of FtsZ from *B. subtilis*). In most cyanobacteria and chloroplast FtsZs, the corresponding amino acids are Ala (A271 and A274 of SyFtsZ). Interestingly, there are several cyanobacterial FtsZs, such as FtsZ from *Prochlorococcus* and *Synechococcus*, where these are still Ser and Thr. Both of these belong to marine picocyanobacteria, whose cell diameter is less than 1 μm. This size is similar to that of bacteria, and much smaller than other cyanobacteria. Also, some slow-growing bacterial FtsZs have only one Thr or Ser. For example, in *M. tuberculosis* FtsZ (MtbFtsZ), the first Thr is changed to Gly, and in *S. pneumoniae* FtsZ (SpnFtsZ), the second Ser is changed to Ala. Previous results suggested that both MtbFtsZ and SpnFtsZ assembled into small bundles. MtbFtsZ slowly assembled into double strands pfs at pH 6.5 (12,31), and SpnFtsZ assembled into mostly double or multiple strands pfs in solution with a high critical concentration around 3-6 μM (32,33).

**Figure 6.**
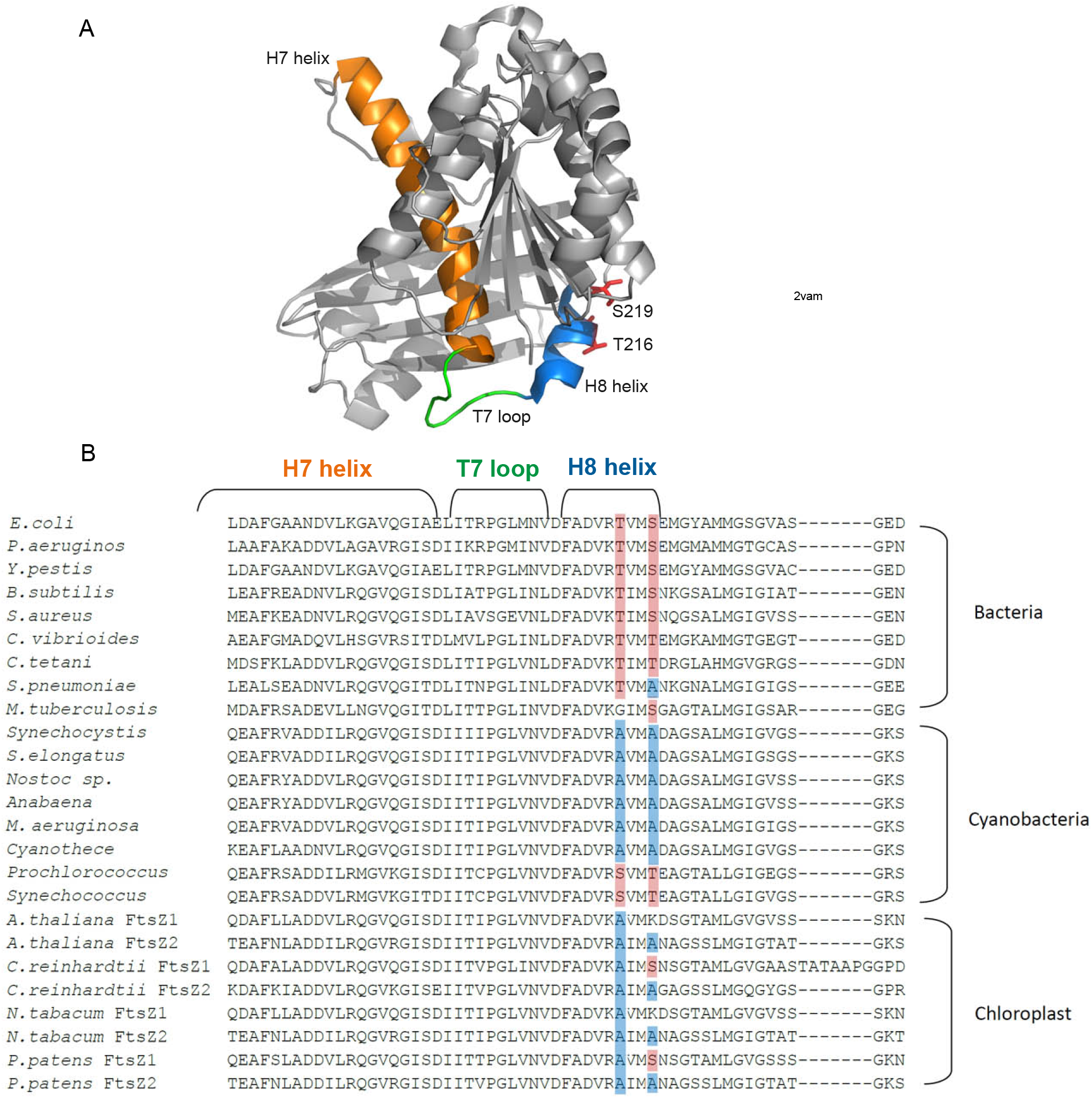
(A) Structure of FtsZ from *B. subtilis* (PDB code 2vam). The H7 helix is shown in orange, T7 loop is green and H8 helix is blue. T216 and S219 are shown as sticks. (B) Sequence alignment of FtsZ with a focus on the conserved region of H7 helix, T7 loop and H8 helix from bacteria, cyanobacteria and chloroplasts. Two amino acids in helix H8 are mostly conserved as Thr and Ser in bacteria (highlighted in red) and mostly Ala in cyanobacteria (highlighted in blue).

To address whether these two amino acids are related to bundle formation, we constructed the double mutant, SyFtsZ-A271T/A274S (SyFtsZ-TS). The GTPase activity of this mutant increased ~80%: from 0.4 to 0.7 GTP per FtsZ per minute in pH 7.5 buffer (Fig. 7A) and from 0.5 to 0.9 GTP per FtsZ per minute in pH 7.2 buffer (Fig. 7B). Fig. 7 (C and D) highlight the difference of their assembly properties measured by the light-scattering at different pH. In pH 7.5 solution, the assembly of mutant SyFtsZ only had the fast initial stage in contrast to the two-stage assembly curves of wild type SyFtsZ assembly (Fig. 7C; compare to Fig. 2B). Negative stain EM confirmed that the assembled pfs of SyFtsZ-TS were mixtures of single pfs and thin straight and curved bundles, including some closed circles (Fig. 7E). There were no large bundles in pH 7.5 buffer for FtsZ up to 15 μM. In pH 7.2 solution, the assembly showed two stages, but they were much slower than wild type (Fig. 7D). The first stage of assembly, which plateaued from 50 to 200 s, showed short pfs and circles typical of this stage (Fig. 7F). The second stage started around 200 s, rose to a much higher light-scattering signal and approached a plateau at 900 s (Fig. 7D). EM showed mostly thick straight bundles and a few circles at pH 7.2 (Fig. 7G), and no toroids-like circle bundles were observed. Evidently, mutant proteins reduced bundles formation.

**Figure 7.**
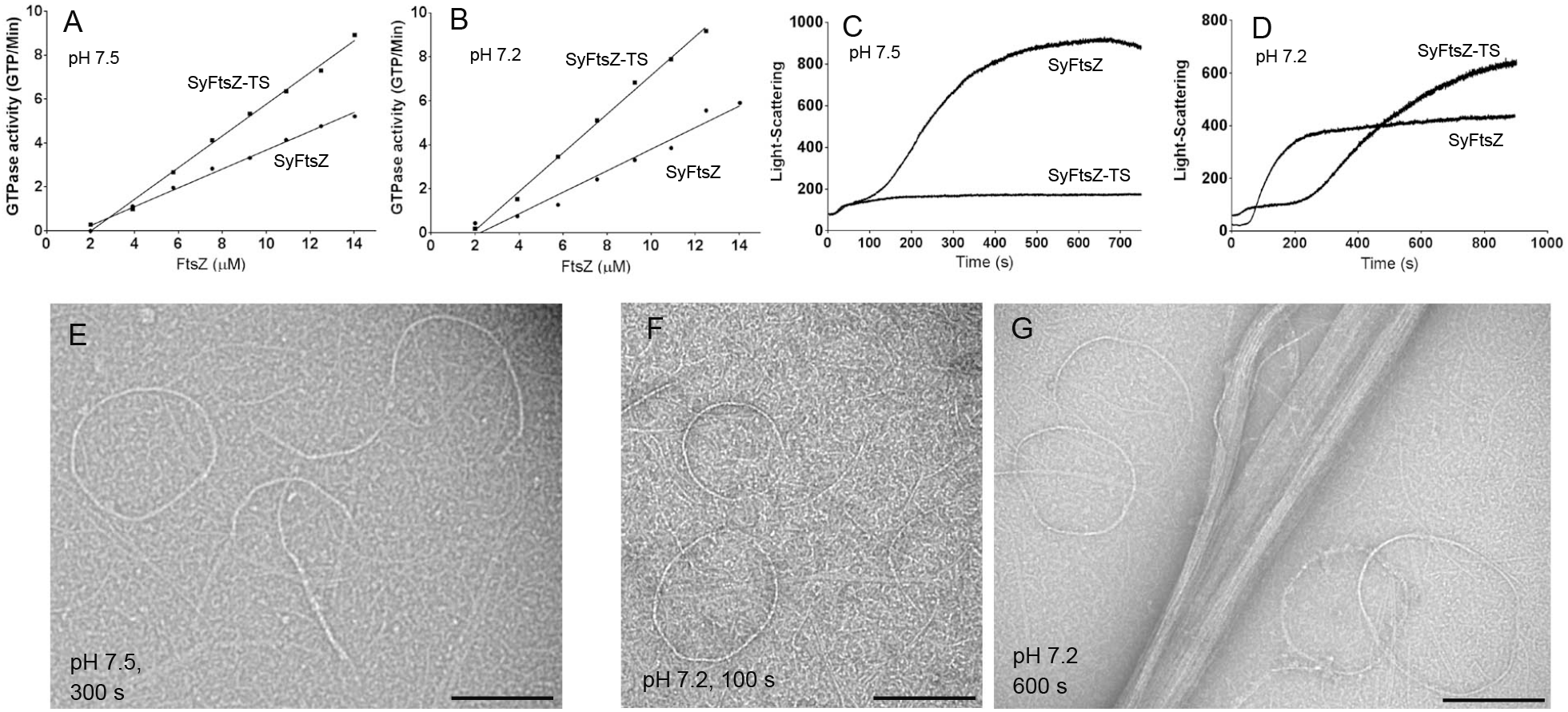
Comparison of assembly properties of wild type SyFtsZ and SyFtsZ-TS (SyFtsZ-A271T/A274S). (A, B) The double mutant increased the GTPase somewhat at both pH 7.5 (A) and pH 7.2 (B). (C) Light-scattering suggests that the SyFtsZ-TS at pH 7.5 assembled only to the first stage, and was missing the large rise in turbidity attributed to the second stage (cf Fig. 2). (D) At pH 7.2 the first stage of assembly was prolonged, but the second stage rise in turbidity began at ~300 s. (E-G) EM confirms that SyFtsZ-TS only assembles into single straight pfs and thin curved bundles at pH 7.5, 300 s (E), and at pH 7.2, 100 s (F). These assemblies are characteristic of stage one (see Fig. 2). At pH 7.2, 600 s large thick bundles characteristic of stage 2 are seen. Bar is 200 nm.

We then tested the effect on EcFtsZ of changing the Thr and Ser to Ala. We constructed two single mutants (EcFtsZ-T215A, EcFtsZ-S218A) and a double mutant EcFtsZ-T215A/S218A. Negative stain EM showed a modest increase in bundles assembled for the mutant EcFtsZ (Fig. 8A-D). These were mostly double pfs or small bundles, and no large bundles were seen. These modest bundling filaments were similar to the assembly of MtbFtsZ (12,31) and SpnFtsZ (32), which had one of the two amino acids changed. Interestingly, GTPase activities of mutant EcFtsZ were only slightly changed (Fig. 8E). The GTPase was reduced from 4.6 to 3.5 GTP per FtsZ per minute for EcFtsZ-S218A, and little changed for EcFtsZT215A and EcFtsZ-T215A/S218A. The critical concentration increased from 1 to around 2.5 μM for EcFtsZ-T215A and EcFtsZ-T215A/S218A.

**Figure 8.**
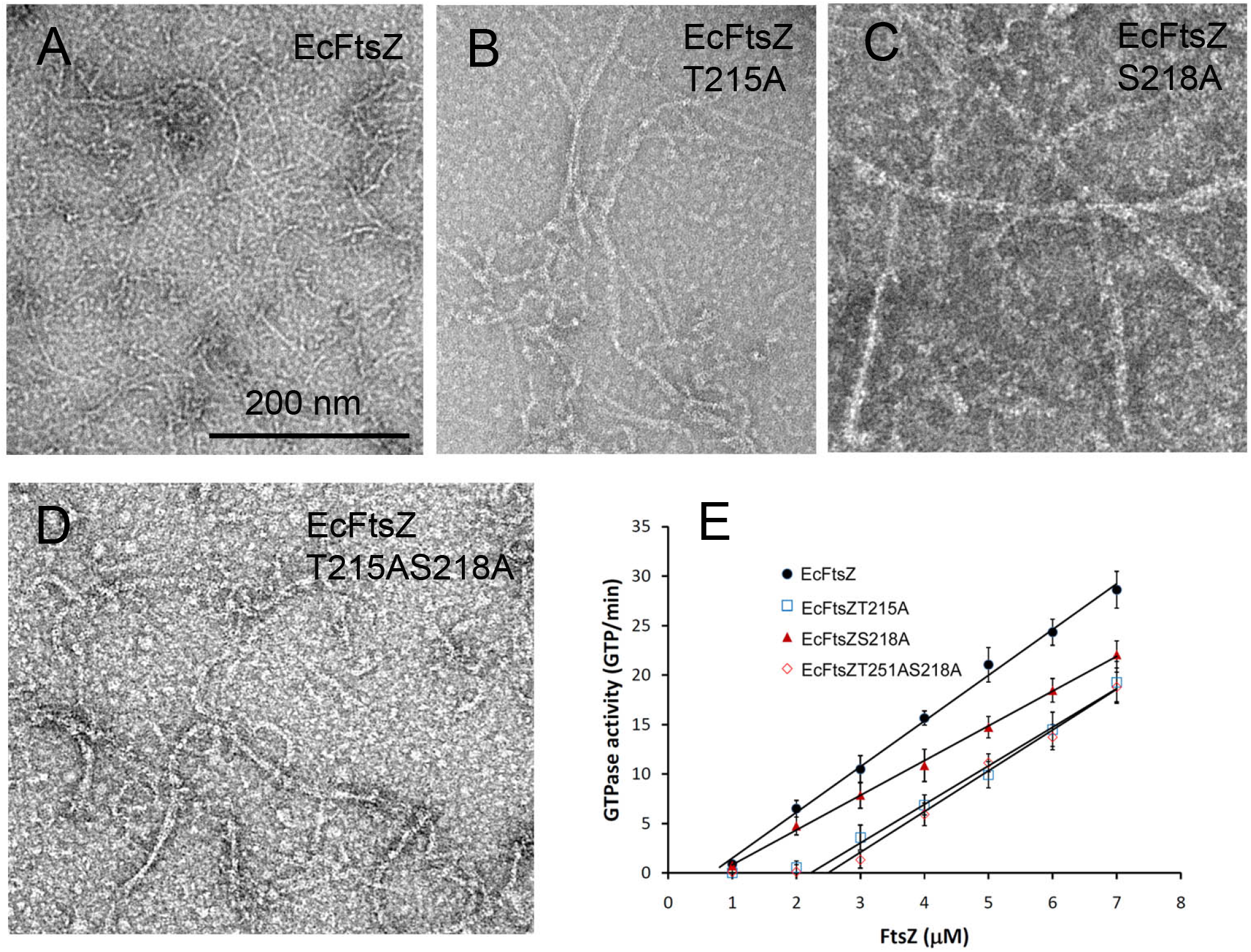
(A-D)Assembly of EcFtsZ with T215 and S218 changed to Ala, singly or together. All mutants showed a modest increase in pf pairs and thin bundles. Bar is 200 nm. (E) GTPase activity showed only small changes for the mutants.

## Discussion

Perhaps the most striking observation in the present work was that SyFtsZ assembled into thick bundles of pfs that could be either straight or curved into a toroid of 200-300 nm diameter. Assembly of similar straight and curved pf bundles was reported previously for EcFtsZ and MtbFtsZ, but only in the presence of crowding agents polyvinyl alcohol or methylcellulose (34,35). A recent study of FtsZ from the cyanobacterium *Anabena* showed a mixture of straight pf bundles and toroids essentially the same as those we show here for SyFtsZ. In that study removal of the 51 N-terminal amino acids (this N-terminal peptide is much longer in cyanobacteria than the 9-15 amino acids in most bacteria) eliminated the toroids but left the thick straight pf bundles. FtsZ from the chloroplasts of *Galdieria sulphuratia*, GsFtsZA and GsFtsZB, also formed thick straight bundles. In this case, the assembly used the globular domains only, truncating the N- and C-terminal extensions.

The thick bundles seem to be a property of cyanobacterial FtsZ and the derivative chloroplast FtsZ. However, the relevance of the bundling for *in vivo* function is not clear. The *in vitro* bundles appear to be three dimensional, but since all FtsZ subunits have the C-terminal peptide that mediates membrane tethering, the *in vivo* bundle may be limited to a two-dimensional ribbon, with all pfs tethered to the membrane. The amount of FtsZ in cyanobacteria has not been determined, but in other bacteria, it is typically present at ~5,000 copies (2). This could make a ribbon of 7 pfs encircling the 1 μm diameter, in *E. coli*. Where only 30% of the FtsZ is in the Z ring, the ribbon would average only 3 pfs wide. There is not enough FtsZ in a single cell to make a large bundle.

We found that bundling was reduced significantly when we mutated A271 to Thr and A274 to Ser, which were the amino acids at the corresponding positions in *E.* coli and other rapidly dividing bacteria. Interestingly, in the FtsZ from marine picocyanobacteria, such as *Prochlorococcus* and marine *Synechococcus*, these amino acids are still Ser and Thr, suggesting that they may reduce the bundling activity. Those marine picocyanobacteria have small sizes with diameters of less than 1 μm. Previous results revealed that marine *Synechococcus* strains and *Prochlorococcus* were sister clades, that was distantly related to other cyanobacteria, including freshwater *Synechococcus* strains, such as *Synechococcus elongates* (36–39). It is consistent with our analysis. One possibility is that the intrinsic bundling activity is useful for the stable Z-ring formation and benefit for the larger diameter of most cyanobacteria and chloroplasts. The biochemical properties of marine picocyanobacterial FtsZ need to be further investigated. Also, we don’t have *in vivo* data to support it; further investigations need to be checked in the future.

The physiological function of the circle bundles of cyanobacterial FtsZ is still unclear. We observed that SyFtsZ assembled into both straight and circle bundles when pH were above 6.8; meanwhile, only straight bundles assembled when pH was 6.5. It was reported that cyanobacterial cell maintained a narrow range of survivable intracellular pH of 7.1-7.3 even when the cell was exposed to low, growth-inhibiting external pH of below 6 (40). It indicates that the FtsZ circles might be useful for its function. Recently, Corrales-Guerrero L et al (41) reported that deletion of N-terminus of *Anabaena* FtsZ, another cyanobacterial FtsZ, only formed straight bundles. These mutant FtsZs might cause the aberrant Z-structures and asymmetric cell division. It indicated that the formation of the medium circles might be necessary for its function, and even it cannot be excluded the possibility that deletion might affect the binding with the other association proteins (41).

The 200-300 nm curvature of the toroids has been designated the intermediate curved conformation (42). Osawa and Erickson have suggested that this intermediate curved conformation, rather than the highly curved miniring, may be responsible for generating the constriction force on the membrane (43). It is not clear what triggers the transformation from straight to intermediate curved. GTP hydrolysis is not required, since intermediate curved pfs can be obtained in non-hydrolysable GMPCPP and by the EcFtsZ mutant D212A (2). Also, Popp et al (34) observed toroids and spiral pf bundles assembled in GMPCPP and in GTP plus EDTA. Previous studies have provided evidence that a pf bundle can transition from straight to curved, in a process called spooling (44,45). A transition from straight to curved pfs has been invoked for treadmilling pf patches (46). Our Fig. 1 provides some of the clearest images of the spooling transition for SyFtsZ.

Another type of curved pfs is defined as highly curved pfs and minirings. The miniring with a diameter around 20-30 nm were first reported when EcFtsZ was assembled onto cationic lipid monolayers without GTP or after GTP hydrolysis (22,47). The miniring structures of EcFtsZ could also be observed in solution (24,25,48), especially following the addition of ZipA (24), a FtsZ associate protein conserved in only γ-proteobacteria. Similar miniring structures were further observed in the studies of *C. crescentus* FtsZ when stabilized by the regulator protein FzlA (49) and *B. subtilis* FtsZ with the addition of ZapA(50). It is concluded that GTP led the assembly of straight pfs or intermediated curved pfs, and GDP favored highly curved pfs assembly. It is possible that these two curved conformations of FtsZ together provide a continuous force to bend the membrane and finish the cell division.

## EXPERIMENTAL PROCEDURES

### Plasmid construction and protein purification

cDNA for SyFtsZ from cyanobacterium *Synechocystis* sp. PCC 6803 was inserted into pET15b using PCR and engineered NdeI/BamHI digestion sites. Mutants SyFtsZ-A271T/A274S (SyFtsZ-TS), EcFtsZ-T215A, EcFtsZ-S218A and EcFtsZ-T215A/S218A were generated by point mutagenesis.

Plasmids containing SyFtsZ or EcFtsZ were transformed into BL21 *E. coli* and purified. Briefly, the soluble His6 proteins were first purified using a Talon column (Clutch Lab, Inc.). After elution with 250 mM imidazole, the samples were concentrated using an Amicon Ultra-15 Centrifugal Filter (Millipore Sigma) and exchanged into LTK50 buffer (50 mM Tris, pH 7.9, 50 mM KCl, 1 mM EDTA, 10% Glycerol). Thrombin was added to 20 units/ml and incubated at room temperature for 2-4 hours to remove the His6 tag. Further purification was over a Resource Q column (GE healthcare) with a linear 50 mM – 1 M KCl gradient in LTK buffer. Purified proteins were stored at -80° C. Protein concentrations were determined by using a BCA assay with a corrected factor for a presumed 70% color ratio (51). Proteins were thawed and dialyzed into HMK buffer (50 mM Hepes, 100 mM KAc, 5 mM MgAc) for assembly at pH 7.5 and 7.2, or MMK buffer (50 mM MES, 100 mM KAc, 5 mM MgAc) for assembly at pH 6.8 and 6.5.

### Electron Microscopy

Negative stain electron microscopy was used to visualize FtsZ polymerization. FtsZ proteins were incubated with GTP or GDP for 2-10 minutes. Then ~10 μl was applied to a carbon-coated copper grid and quickly withdrawn. Grids were immediately stained with 2% uranyl acetate, and specimens were imaged with a Philips 420 electron microscope.

### Light-scattering measurement

The kinetics of FtsZ assembly was measured by a light-scattering assay using a Shimadzu RF-5301 PC spectrofluorometer, with both excitation and emission at 350 nm as described previously (20,52). The light-scattering assay is especially sensitive to the assembly of large bundles. Each measurement was repeated two or three times, with consistent results.

### Sedimentation assay

Assembly of SyFtsZ was also assayed by sedimentation. 5 μM or 10 μM SyFtsZ was incubated with or without 1 mM GTP in HMK buffer, pH 7.5 or MMK buffer, pH 6.5 at room temperature for 10 min and centrifuged at 50,000 rpm or 90,000 rpm for 30 min at 25 °C in a Beckman TLA100 rotor. The supernatant was carefully removed, and the pellet was resuspended in the same volume of buffer. The protein in the pellet and supernatant was assayed by 10% SDS-PAGE. The ratio of supernatant and pellet was measured using ImageJ software.

### GTPase measurement

The GTPase activity of FtsZ was measured using a regenerative coupled GTPase assay as described previously (52,53). Our assay mixture included 1 mM phosphoenolpyruvate, 0.8 mM NADH, 20 units/ml pyruvate kinase and lactate dehydrogenase (Sigma–Aldrich), and 0.5 mM GTP. After GTP hydrolysis by FtsZ, GDP in solution is rapidly regenerated into GTP, in a reaction that consumes one NADH per GDP. The NADH concentration was measured by the absorption at 340 nm in a Shimadzu UV-2401PC spectrophotometer, using the extinction coefficient 6,220 M-1 cm-1. The absorbance showed a linear decrease over time, giving the hydrolysis rate for each FtsZ concentration. These rates were plotted versus FtsZ concentration, and the overall hydrolysis rate in GTP per minute per FtsZ was the slope of the line above the critical concentration. Each assay was repeated two or three times.

## Acknowledgments

This work was supported by First-class University and Academic programs of Northwest University (to Y.C.). I thank Dr. Harold Erickson, Duke University, for providing laboratory and electron microscopy resources for some of this work, and for helpful comments on the manuscript.

## Author Contributions

YC and PW conceived the project; YC, NW, and LB did most of the experimental work and interpretation; XM, CSC and MR contributed experimental work and interpretation; YC and PW wrote the manuscript with contributions from all authors. All authors reviewed the results and approved the final version of the manuscript.

## FOOTNOTES

### Abbreviations

SyFtsZ: FtsZ from the cyanobacterium *Synechocystis* sp. PCC 6803
EcFtsZ: FtsZ from *Escherichia coli*
pf: protofilament
pfs: protofilaments
EM: electron microscopy
SyFtsZ-TS: double mutant SyFtsZ-A271T/A274S

